# Testing models of cell cortex wave generation by Rho GTPases

**DOI:** 10.1101/2024.04.29.591685

**Authors:** Dom Chomchai, Marcin Leda, Adriana Golding, George von Dassow, William M. Bement, Andrew B. Goryachev

**Author notes:** Equal contributions (Dom Chomchai, Marcin Leda). Equal contributions (George von Dassow, William M. Bement, Andrew B. Goryachev).

## Abstract

The Rho GTPases pattern the cell cortex in a variety of fundamental cell-morphogenetic processes including division, wound repair, and locomotion. It has recently become apparent that this patterning arises from the ability of the Rho GTPases to self-organize into static and migrating spots, contractile pulses, and propagating waves in cells from yeasts to mammals^1^. These self-organizing Rho GTPase patterns have been explained by a variety of theoretical models which require multiple interacting positive and negative feedback loops. However, it is often difficult, if not impossible, to discriminate between different models simply because the available experimental data do not simultaneously capture the dynamics of multiple molecular concentrations and biomechanical variables at fine spatial and temporal resolution. Specifically, most studies typically provide either the total Rho GTPase signal or the Rho GTPase activity as reported by various sensors, but not both. Therefore, it remains largely unknown how membrane accumulation of Rho GTPases (i.e., Rho membrane enrichment) is related to Rho activity. Here we dissect the dynamics of RhoA by simultaneously imaging both total RhoA and active RhoA in the regime of acute cortical excitability^2^, characterized by pronounced waves of Rho activity and F-actin polymerization^3-5^. We find that within nascent waves, accumulation of active RhoA precedes that of total RhoA, and we exploit this finding to distinguish between two popular theoretical models previously used to explain propagating cortical Rho waves.

## Results and discussion

### Rho GTPase activity rise precedes accumulation of total Rho in waves

To provide a detailed analysis of spatio-temporal dynamics of RhoA in a dynamic cortical patterning process, we amplified cytokinetic waves in starfish oocytes undergoing meiosis via ectopic expression of the cytokinetic Rho GEF Ect2^3^ or induced such waves in immature frog oocytes via ectopic expression of Ect2 and the cytokinetic Rho GAP RGA-3/4^4^. Such waves mimic those in the cytokinetic apparatus^6^ but have the virtue of continuing for many minutes or hours and encompassing the majority, if not the entirety, of the cell cortex, permitting a detailed record of their behavior to be captured. In contrast, the furrow waves are normally restricted to anaphase and quickly become inaccessible to high resolution imaging as the furrow ingresses. As a result, amplified and induced waves have been used to probe the network dynamics of cytokinetic Rho GTPase signaling^3,7^, wiring of the underlying signaling networks^4^, and relationships between cell shape, wave propagation and signaling hierarchies^8,9^. More broadly, propagating waves offer distinct advantages over other cortical states that lack the dynamicity of waves. Firstly, waves present as readily distinguishable alternating maxima and minima of fluorescence intensity moving in a periodic pattern. Thus, standard image analysis methods can be readily applied to extract wave velocity, spatial wavelength and temporal period. These constitute a quantitative signature of the wave pattern, which can be compared to proposed theoretical models^4,10^. Secondly, since wave propagation consists of periodically repeating cycles of biochemical reactions and molecule translocations between the plasma membrane and the cytoplasm, time series of fluorescence signals carry valuable mechanistic information. For example, they reveal the temporal sequence of various processes, which may suggest causal relationships between them. Such data informs the construction of plausible mechanistic models. Further, specific time delays, measured, for example, between the concentration maxima of the key molecular players, provide information necessary to quantitatively constrain the values of parameters to be associated with the model interactions.

Since the membrane-bound pool of RhoA consists of dynamically interconverting active and inactive GTPase states, two independent fluorescence signals representative of the total RhoA and active, GTP-bound RhoA, are required for the complete characterization of RhoA dynamics at the cell cortex. To visualize total RhoA, we employed previously characterized, internally-tagged frog RhoA^11^ (IT-RhoA) and developed the equivalent echinoderm version for starfish. Both probes localize to the cytokinetic furrow^11^ (Figure S1A) and their furrow localization is amplified by co-expression of Ect2 (data not shown). The merits of internal fluorescent protein tagging of Rho GTPases have been pointed out previously for both yeast^12-14^ and vertebrates^11,15^. To visualize RhoA-GTP, we employed mCherry-rGBD, which has been used previously in both starfish and frog to track active RhoA^3,9,16,17^. Starfish IT-RhoA produced a robust fluorescence signal in Ect2 amplified waves in starfish oocytes (Figure S1B). Interestingly, direct comparison revealed that IT-RhoA waves follow RhoA-GTP waves with a delay, which is evident in *en face* movies, still images, and kymographs as a color shift (Figure 1A (i), Movie S1). This delay is also evident from intensity plots of the raw data (Figure S1C). To quantify the delay, wave amplitude was normalized to lie between 0 (minimum) and 1 (maximum), which allows for more precise quantification of phase differences against a background of data variability (see Methods). (We emphasize that normalized plots do not preserve the relationship between the absolute levels, as active RhoA always represents some fraction of the total RhoA). Correlation analysis of the normalized data from a representative cell showed that IT-RhoA waves followed RhoA-GTP waves with a 10.91 ± 0.69 sec delay (Figure 1A (ii)). Quantification of the normalized data from multiple cells revealed an average of 9.3 ± 2.4 sec (n=20 cells from 3 experiments; Figure 1A (ii)) corresponding to an oscillation phase shift of 9.5% (period = 97 ± 18.7 sec).

**Figure 1.**
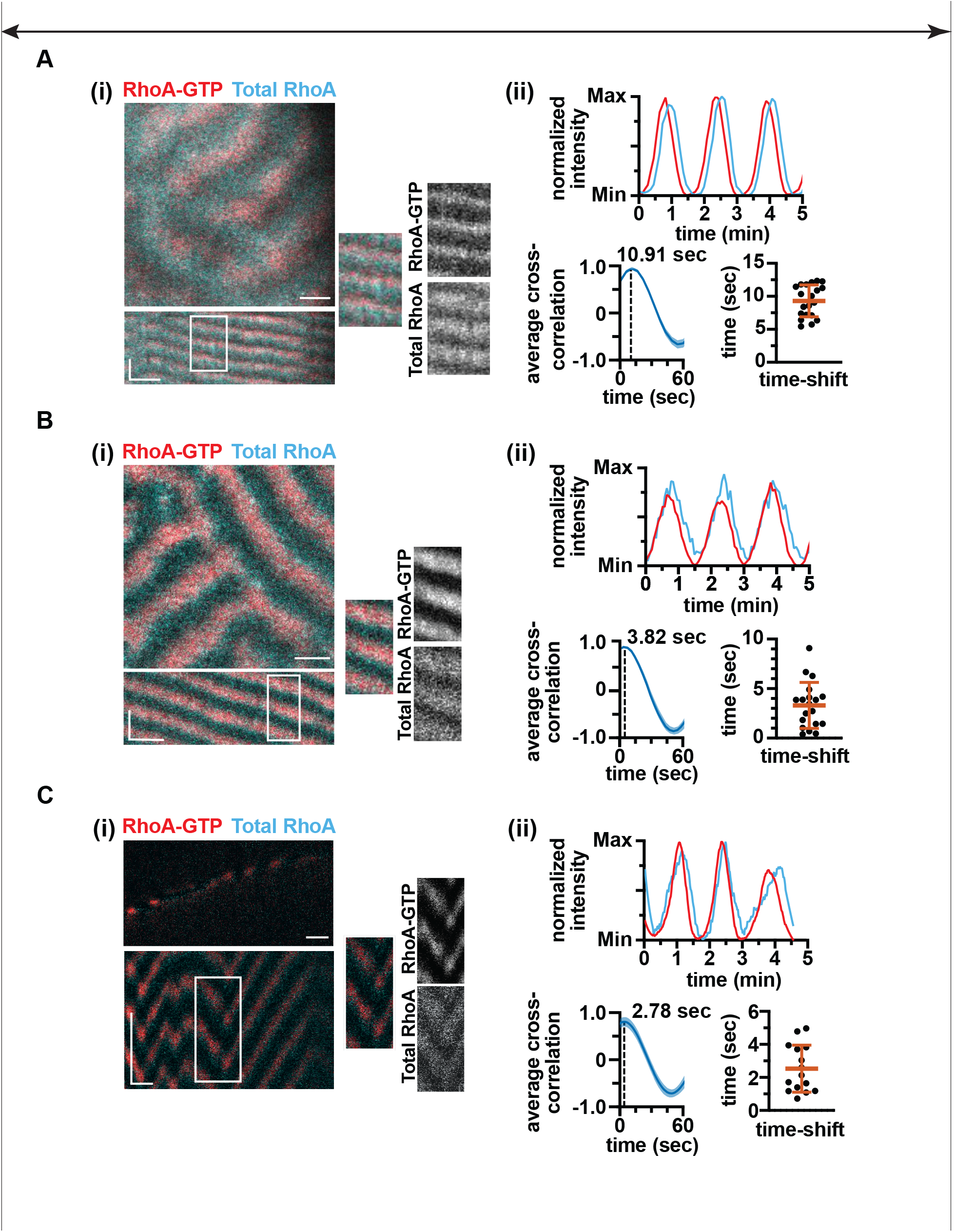
RhoA activity leads RhoA membrane enrichment in starfish and frog oocyte cortical excitability. (A-C(i)) Representative still images and corresponding kymographs of RhoA-GTP (red) and Total RhoA (cyan) waves from a starfish oocyte *en face* movie (A), a frog immature oocyte *en face* movie (B), and a frog immature oocyte medial section difference movie (C). (ii) Quantification for the corresponding movies in A-C. Top: Representative normalized quantitative wave profiles of RhoA-GTP (red) and Total RhoA (cyan) waves from the corresponding movie in A-C demonstrate that Rho-GTP leads total Rho enrichment. Bottom left: Average cross-correlation of the corresponding movies in A-C. Shaded area – standard deviation (SD). Vertical dashed line – mean time-shift between the Rho-GTP/Total Rho signals. (A: 10.91 ± 0.69 sec; B: 3.82 ± 1.10 sec; C: 4.99 ± 2.86 sec; ±SD). Bottom right: Overall mean time shift between Rho-GTP and Total Rho signals. Each point represents the time shift measurement of an individual cell under the same conditions as A-C (A: 9.30 ± 2.41 sec, n=20; B: 3.30 ± 2.32 sec, n=19; C: 2.53 ± 1.4 sec, n=15; ±SD; p-value <0.0001 for each; two-tailed one-sample t-test). Horizontal scale bars: 10 μm. Vertical scale bars: 2 min.

In frog, IT-RhoA also produced a robust fluorescence signal in induced waves (Figure S1D). However, the shift between IT-RhoA and RhoA-GTP signals, while still detectable in *en face* movies, their kymographs, and the normalized time series (Figures 1B, S1E and Movie S2), was much more subtle than in starfish, with a shift of 3.82 ± 1.10 sec. Quantification of multiple cells revealed an average shift of 3.30 ± 2.32 sec (n=19 cells from 5 experiments), which was less than the interval of image acquisition. We therefore adopted single optical plane imaging in a medial section (Figure 1C(i), Movie S3), which shortened the sampling interval from ∼4 sec to ∼1 sec. Consistent with the *en face* analysis, this revealed a small but significant delay of 2.78 ± 1.41 sec from a representative cell and an average of 2.53 ± 1.4 sec; n=15 cells from 4 experiments (Figure 1C(ii)). This constitutes only a 3.6% oscillation phase shift (period = 70 ± 17.0 sec). However, in both starfish and frog, there must be some inherent delay between the activation of the GTPase and binding of the probe, implying that the measurements actually underestimate the delay between the active and total RhoA.

Importantly, these results clearly demonstrate that the waves of Rho GTPase activity reported by us and others in the oocytes and embryos of frogs^3,4^ and echinoderms^3,4,9,18^ are not simply waves of local GTPase activation-inactivation against a background of constant total plasma membrane-associated GTPase. Rather, they show for the first time that the waves comprise changes in both RhoA activity and total plasma membrane RhoA levels. Further, the delay between the peaks of RhoA activity and total RhoA unambiguously indicates that the change in GTPase activity precedes the change in its total quantity, thus suggesting a causal relationship between the two.

### Testing theoretical models of RhoA wave generation

We next asked if our experimental results could shed additional light on the molecular mechanisms of Rho GTPase dynamics and, consequently, wave generation. Theoretically, two distinct types of minimal mechanistic models – activator-inhibitor and activator-depleted substrate (ADS) – could explain propagation of waves in the cortex^19^. Both models assume positive feedback in which the activator stimulates its own activation or production, but they differ in how their negative feedback is implemented: In activator-inhibitor models, negative feedback arises from an inhibitor whose accumulation or activity is stimulated by the activator. This inhibitor inactivates or antagonizes the production of the activator. As a result, a wave of activator is “chased” by a wave of inhibitor. In ADS models, negative feedback arises from the consumption of some substrate essential for the activator production. As a result, the wave of activator is chased by a wave of depletion of the substrate in question. In the specific context of dynamic pattern formation by Rho GTPases, RhoA-GTP is the activator and engages in positive feedback by directly or indirectly stimulating a Rho GEF^1,20,21^. In activator-inhibitor models for the GTPases, the inhibitor is surmised to be a GTPase activating protein (GAP) that is a downstream target of active GTPase^1,22-25^. These models are appealing based on demonstrations of actin filaments (F-actin) serving as a prototypical inhibitor in a number of systems^3,26-28^, at least in part based on their ability to associate with a Rho GAP that binds reversibly to F-actin and inactivates RhoA^4,27-29^. ADS-type models for the Rho GTPases are appealing because they have the potential to apply to any GTPase that undergoes nucleotide cycling. In the ADS models, RhoA-GDP is considered the essential substrate subject to depletion. Indeed, regardless of any additional interactions, all molecular mechanisms of wave generation based on small GTPases incorporate the activator-depleted substrate network motif (common elements on Figures 2A,B), simply because the active GTPase is generated from the inactive GTPase.

**Figure 2.**
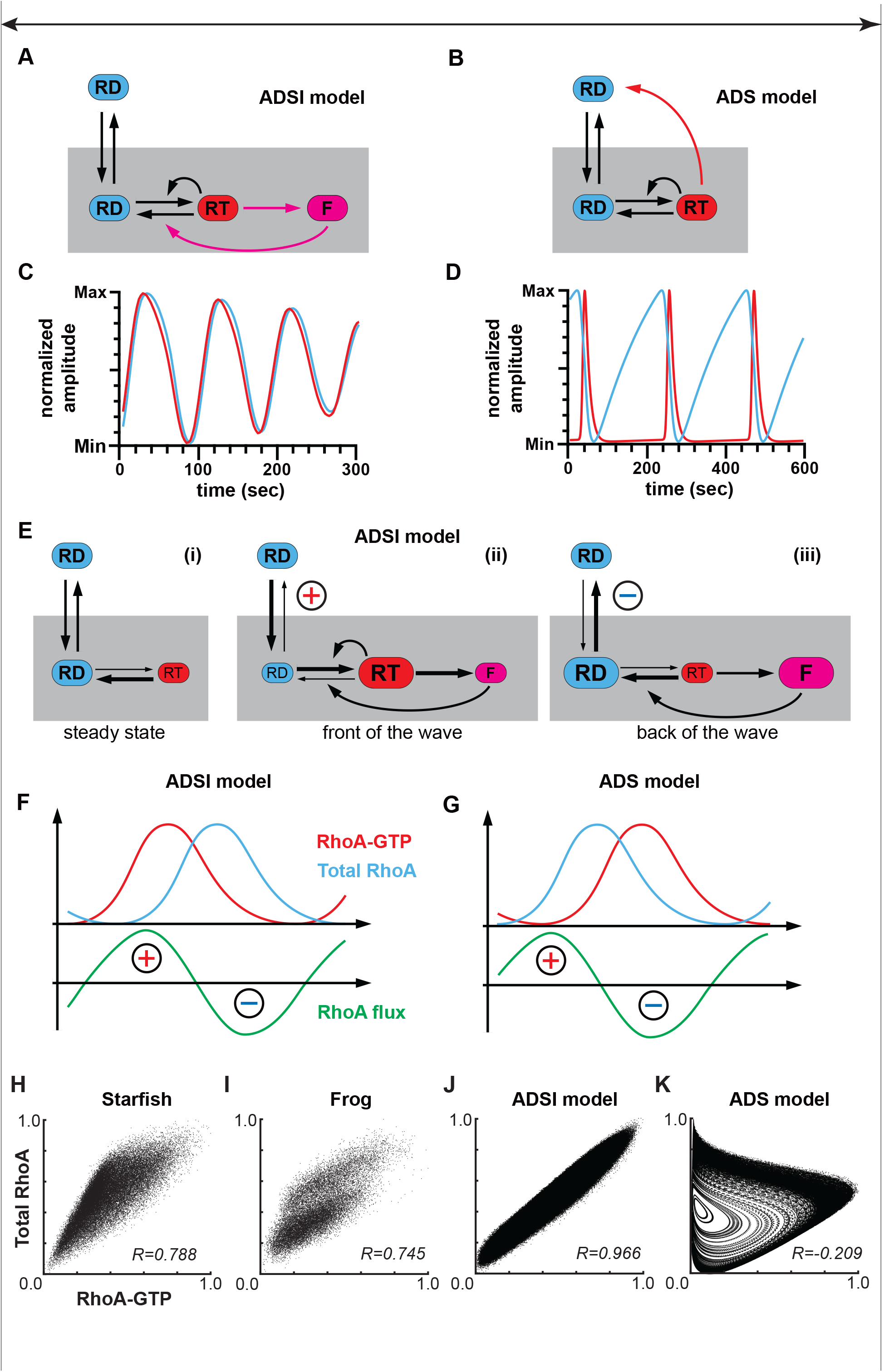
Comparison of the ADSI and ADS models of wave dynamics. (A,B) Schematic diagrams of reaction networks. RT – RhoA-GTP, RD – RhoA-GDP, F – inhibitor. Shaded area represents membrane. (C,D) Representative time series of the spatially-distributed ADSI (C) and ADS (D) models. Red, RhoA-GTP, cyan, total RhoA. (E) Schematic diagrams of Rho dynamics according to the ADSI model. Sizes of labels and arrows represent relative magnitudes of molecular pools and fluxes, respectively. (F,G) Schematic diagrams comparing profiles of RhoA-GTP, total RhoA and RhoA flux predicted by ADSI (F) and ADS (G) models. (H-K) Normalized dynamics of specified on the figure systems shown as scatter plots. (H) Averaged pixel values from a starfish movie; (I) same for a frog medial section movie; (J) data from the spatially distributed simulation of the ADSI model; (K) data from the spatially distributed simulation of the ADS model.

An explicit inhibitor can be added to the ADS model as shown in Figure 2A and the resulting molecular mechanism is then described by a three-variable activator-depleted substrate-inhibitor (ADSI) model. This circuit consists of the activator (RT)-depleted substrate (RD) and the activator (RT)-inhibitor (F) submodules linked by a common activator, a layout often inferred in systems exhibiting complex oscillatory dynamics^30-32^. However, in the framework of the ADSI model^3,4^, depletion of inactive RhoA does not play a key role in the generation of waves, which are instead produced by the action of the inhibitor, F (Figure 2A). Importantly, analysis of the ADSI model shows that, under the biologically realistic parameter combinations conducive to wave generation, the maximum of the RhoA-GTP concentration in waves always precedes that of the total RhoA concentration (Figure 2C) in agreement with our experimental data.

This model prediction can be understood within the framework of the theoretical concept of GTPase flux^1,33-35^ which is based on the premise that RhoA-GDP participates in rapid membrane-cytoplasmic exchange, while RhoA-GTP is largely membrane-bound until inactivated by GAPs. This conjecture is strongly supported by the observation that Rho GDP dissociation inhibitors (Rho GDIs), which solubilize Rho GTPases, have a higher affinity for Rho-GDP than Rho-GTP^22,36,37^, the demonstration that Rho GTPases in the cytoplasm are predominantly in the GDP-bound form^38,39^, and the observation that Rho-GTP is enriched on membranes^23,40^. In a spatially homogeneous steady state (i.e., in the absence of any spatial patterns), the cytoplasmic and membrane pools of inactive RhoA are in equilibrium resulting in a zero net RhoA flux (Figure 2E(i)). In a cortex exhibiting waves, activation of RhoA by the membrane-localized GEFs at the front of the wave drives the system out of equilibrium, leading to the local depletion of the membrane-bound pool of RhoA-GDP. This depletion is rapidly compensated for by the translocation of RhoA-GDP from the cytoplasm into the depleted region of the membrane (positive Rho flux, Figure 2E(ii)). The newly deposited RhoA-GDP is subsequently activated on the membrane inducing more RhoA-GDP depletion and thus more compensatory RhoA-GDP deposition. This positive feedback loop of RhoA activation and accumulation proceeds until it is halted and then reversed by the negative feedback provided by the inhibitor. At the back of the wave, inactivation of RhoA causes the local surplus of membrane-bound RhoA-GDP that is recycled back to the cytoplasm (negative Rho flux, Figure 2E(iii)). Thus, the system generates a GTP-powered molecular current, in which the maximum of GTPase activity approximately coincides with the maximum of the GTPase positive flux (Figure 2F). While the GTPase flux cannot be directly measured experimentally, the time delay observed here between the maxima of RhoA activity and membrane enrichment of total Rho strongly supports the ADSI model.

What does the ADS model predict? Here, an explicit inhibitor is not required because negative feedback is implicitly embedded in the form of depletion of inactive RhoA caused by its local rapid activation (*RD* → *RT*). The ADS-type models accounting for the RhoA membrane-cytoplasmic shuttling and nucleotide cycling (black arrows, Figures 2A,B) were originally developed to explain the spontaneous formation of stationary spots or clusters of GTPase activity in cell migration and yeast budding^1,33,41,42^. However, to account for propagating waves of GTPase activity, the ADS model requires additional mechanistic assumptions. Specifically, mathematical analysis shows that to be able to explain waves, the ADS model strictly requires that the membrane association lifetime of RhoA-GTP must be considerably shorter than that of the inactive GDP-bound form (see Methods and Figure S2). Thus, a reaction of simultaneous inactivation and membrane detachment of active RhoA has been recently suggested^9,43^ (red arrow, Figure 2B). One period of wave dynamics, according to this model (Figure 2B), involves progressive membrane accumulation of inactive Rho followed by its rapid autocatalytic activation (Figure 2D). The RhoA-GTP thus produced is then simultaneously inactivated and removed from the membrane (red arrow, Figure 2B). Together with the depletion of RhoA-GDP from the membrane, this reaction first reverses Rho activation and then fully extinguishes the spike of RhoA activity. This model belongs to the general class of so-called accumulate-and-fire oscillators^44^ known to describe pH oscillations in chemistry^45^. Importantly, by forcing unbinding of RhoA-GTP from the membrane, it induces local depletion of the total membrane RhoA, and the new cycle of activity starts with the replenishment of the membrane pool of inactive RhoA from the cytoplasm (Figure 2D). Thus, the ADS model posits that the maximum of RhoA activity coincides with the maximum of the negative flux of RhoA, i.e., it marks the area of the membrane with the fastest return of Rho to the cytoplasm (Figure 2G). Analysis of the ADS model shows that, as a consequence, the maximum of the total RhoA precedes the maximum of Rho activity (Figures 2D,G), in contrast to the experimental results. This prediction is independent of the specific choice of parameter values and holds true for the ADS model shown in Figure 2B in general (see Methods).

Thus, the sign of the phase shift between the maxima of total RhoA and active RhoA qualitatively separates the two theoretical models of wave generation. We therefore conclude that our data shown in Figure 1 argues strongly in favor of the ADSI model and falsifies the ADS model. As shown in Figure S2C, this conclusion is robust to the unknown time delay between the true maximum of the GTPase activity and that of the fluorescent signal of the probe for active Rho used in our experiments. Indeed, due to this delay, the true maximum of Rho activity is even further ahead of the maximum of total Rho than the measured in experiment maximum of the reporter accumulation (top, Figure S2C). At the same time, if the ADS model held true, the experimentally measured maximum of the activity probe would be even further behind the maximum of total RhoA than the predicted by the ADS model true maximum of RhoA activity (bottom, Figure S2C).

In practice, the calculation of the time delay between the maxima of two signals could be challenging given the noisy data obtained in small cells with insufficient frequency of imaging. A more intuitive approach is demonstrated in Figure 2H,I, where the normalized and spatially averaged intensities of imaging pixels are shown as scatter plots. Comparison of the plots shows that the clouds of data points in both starfish (Figure 2H) and frog (Figure 2I) experiments exhibit high positive correlation, in agreement with the prediction of the ADSI model (Figure 2J). On the contrary, the ADS model predicts a distinct distribution of data points with small negative correlation (Figure 2K).

We found that waves of GTPase activity are unambiguously associated with the waves of GTPase enrichment, consistent with recent results from cell repair^11^ and yeast polarization^13^. Furthermore, in both frogs and starfish, Rho activity waves precede waves of Rho enrichment, a finding that excludes the ADS model, which predicts the opposite result. Thus, the benefit of the ADS model--its reduced complexity due to the lack of the third variable, the inhibitor--is outweighed by its restrictive requirement that the membrane lifetime of active GTPase must be much shorter than the lifetime of its inactive form. These observations directly support the existence of explicit inhibitory molecular complexes that are induced by the activity of GTPases and eventually extinguish it. A network motif in which negative feedback is mediated by a GAP bound to a polymeric network generated downstream of the activity of a small GTPase is likely broadly conserved^1,4^. For example, binding of the Cdc42 GAP Bem2 to the septin scaffold was shown to provide negative feedback to Cdc42 activity in the context of budding yeast polarization^46^. The results presented here also bring into sharper focus the notion of flux of Rho between the cytoplasm and the plasma membrane as a result of the combined action of positive and negative feedback loops, an idea previously only tested for Cdc42 in budding and fission yeast^47-49^. Our results suggest that, in the simplified system exploited here, the molecular circuit established by the expression of the GEF Ect2 and the GAP RGA-3/4 is sufficient to drive such flux. Given the recent demonstrations of close collaboration between Rho GEFs and GAPs in a variety of model systems and cellular contexts^28,50,51^, these findings indicate that Rho flux is likely to be a basic feature of Rho GTPases in cell morphogenetic activity.

## Figure legends

Figure S1. IT-RhoA produces robust waves in starfish and frog oocytes. (A) Representative images of starfish oocytes exogenously expressing Ect2 shows IT-RhoA accumulation in polar body extrusion during Meiosis-II (left) and in the cortical furrow during mitosis (right). (B) Representative image and kymograph of a starfish oocyte exogenously expressing Ect2 produces robust, amplified waves of IT-RhoA. (C) Representative wave profiles depicting the raw signals of RhoA-GTP (red) and IT-RhoA (cyan) extracted from a starfish oocyte exhibiting amplified cortical excitability. (D) Representative image and kymograph of a frog immature oocyte exogenously overexpressing Ect2^dNLS^ and RGA-3/4 displays distinct waves of IT-RhoA. E) Mean active Rho/Total Rho time-shift of individual boxes in starfish oocyte *en face* movies (left, 12.12 ± 1.47 sec; ±SD; p-value <0.0001, two-tailed one-sample t-test) and frog immature oocyte medial section movies (right, 3.39 ± 1.01SD sec: ±SD; p-value <0.0001, two-tailed one-sample t-test) using an independent MATLAB script^10^. Horizontal scale bars: 20 μm; Vertical scale bar: 2 min.

Figure S2. ADS model analysis. (A) Reaction diagram of the ADS model with kinetic rate constants. (B) Phase portrait of the ADS model with phase flow direction indicated on the limit cycle (see Methods for more details on A and B). (C) Comparison of the model predictions and experiment. IT-RhoA, cyan; signal of the activity reporter, red; true unknown profile of RhoA-GTP, red dash.

Movie S1. Representative *en face* movie of RhoA-GTP (red) and Total RhoA (cyan) waves from a starfish oocyte exhibiting amplified cortical excitability.

Movie S2. Representative *en face* movie of RhoA-GTP (red) and Total RhoA (cyan) waves from a frog oocyte exhibiting induced cortical excitability.

Movie S3. Representative medial section movie of RhoA-GTP (red) and Total RhoA (cyan) waves from a frog oocyte exhibiting induced cortical excitability.

## Acknowledgements

ABG acknowledges financial support from the Leverhulme Trust (RPG-2020-220) and from the Biotechnology and Biological Sciences Research Council (BB/W013614/1).

WMB acknowledges financial support from the National Institutes of Health (RO1GM052932).

The authors declare no competing financial interests.

## Method Details

### Oocyte Preparation

Adult *Patiria miniata* (bat stars) were housed in natural seawater tanks with aeration at 11°-14°C, at the Oregon Institute of Marine Biology. Animals were fed with minced, cooked shrimp and locally collected mussels. Oocytes were obtained from ovary fragments, after transfer to Ca^2+^-free artificial seawater. Individual oocytes were kept at 12°C and rinsed several times to remove follicles, and then transferred to filtered sea water at 12°C until microinjection.

Chunks of ovary were collected from adult *Xenopus laevis* females and stored in 1x Barth’s solution (87.4 mM NaCl, 1 mM KCl, 2.4 mM NaHCO3, 0.82 mM MgSO4, 0.6 mM NaNO3, 0.7 mM CaCl2, and 10 mM HEPES at pH 7.6). Oocytes were treated with collagenase for 1 hr at 16°C, and then rinsed extensively with 1x Barth’s solution before recovering overnight at 16°C. Stage VI oocytes were selected and manually defolliculated before injection.

### Constructs and mRNA

All constructs are contained within the pCS2+ vector. The GFP- and mCherry-rGBD used in both starfish and frog experiments were generated by fusing the Rho binding domain of Rhoketin with a fluorescent protein^16^. For frog IT-GFP-RhoA, the construct was created by inserting a GFP protein into an exposed loop of the RhoA protein^11^. A starfish version of IT-GFP-RhoA was developed using a similar approach, where a GFP was introduced within an external loop of the *P. miniata* RhoA. The GDI utilized in this study was previously detailed in Golding et al., 2019. The Ect2^dNLS^ and RGA-3/4 constructs employed for inducing cortical excitability in frogs were described elsewhere^4^. For starfish experiments, we utilized the previously described echinoderm Ect2 construct^3^.

Prior to mRNA synthesis, all plasmids were linearized downstream of the open reading frame. mRNA synthesis was then carried out using the mMessage Machine SP6 kit (Ambion, #AM1340). For starfish mRNA, polyadenylation was performed using the Poly(A) Tailing kit (Ambion, #AM1350). mRNA was subsequently purified using the RNeasy Kit (Qiagen, #74104). Size and quality of mRNA was confirmed via gel electrophoresis, and concentration was calculated against a GFP standard curve.

### Microinjection

Starfish oocytes were sheared with a narrow capillary to remove mucus and arranged in rows on coverslip-bottomed dishes (MatTek) that were pre-rinsed with 1% protamine sulphate for 30 sec. Oocytes were then pressure-injected using capillary glass needles using a Dagan Instruments injector and an Narishige oil-hydraulic micromanipulator. An injection volume corresponding to 1–2% of the total oocyte volume was used and, after injection with mRNA, oocytes were incubated overnight at 12–14°C. The injection needle concentration of IT-GFP-RhoA, Ect2, and rGBD mRNA were 25-50 ng/μL, 50-80 ng/μL, and 50-100 ng/μL, respectively.

Frog oocyte microinjections were performed using a Warner Instruments PLI-100 microinjector and a manual Narishige micromanipulator. Needles were pulled from capillary tubes and calibrated to inject 40 nL of mRNA. Cells were injected in a mesh-bottomed Petri dish containing 1× Barth’s solution. For IT-GFP-RhoA, GDI, and rGBD, mRNA was injected the day before imaging and incubated overnight at 16°C. The injection needle concentration of IT-GFP-RhoA, GDI, and rGBD mRNA were 250 ng/μL, 125 ng/μL, and 100 ng/μL, respectively. For Ect2^dNLS^ and RGA-3/4, the mRNA was injected the morning of imaging, and the cells were incubated at 23°C for >3-4 hours before imaging to allow for cortical wave generation. The injection needle concentration of Ect2^dNLS^ and RGA-3/4 mRNA were 200 ng/μL and 166 ng/μL, respectively.

### Image Acquisition

Starfish oocytes were screened for mRNA expression (as judged with a fluorescent dissecting scope) and small groups were selected for imaging. Oocyte maturation was induced by addition of 1-methyladenine to ∼10^−5^ M. Oocytes were imaged in chambers made by placing 22x30 mm #1.5 coverslips on ∼1 cm lines of vacuum grease drawn on 75x25 mm glass slides with toothpicks. Imaging was conducted on an inverted Olympus FluoView 1000 laser-scanning confocal microscope using a 1.15-NA 40× water-immersion objective. Temperature was maintained at 16–18°C by room air conditioning; these temperatures are well within the range tolerated by *P. miniata*.

Frog oocytes were mounted onto glass slides within a ∼1x1 cm area of vacuum grease to accommodate their size and were covered with #1.5 coverslips in 1x Barth’s solution. Imaging was conducted using a Prairie View Swept Field Confocal (SFC) system mounted on a Nikon Eclipse Ti base (Bruker), employing a 60x 1.4-NA oil immersion objective. The microscope was controlled by Prairie View software (Bruker). Imaging was conducted at room temperature. *En face* imaging was performed by acquiring time-series data with 5 steps of 1μ*m* each. For medial section imaging, images were captured in a single optical plane.

### Image Processing

Image processing was performed using ImageJ/FIJI^52^. Multi-stack *en face* images were max projected, then subsequently divided by the sum of pixel intensities to mitigate vertical artifacts stemming from the SFC microscope. Difference movies were generated in Fiji through the process of subtracting the signal from n frames following a frame of interest from each corresponding frame. Kymographs were generated using Fiji’s reslice function, with a 1-pixel-wide line across the field of view for en-face imaging or along the cell edge for medial section imaging. All images and figures were compiled in Adobe Illustrator.

### Image Analysis

The analyses depicted in Figure 1, including time-shift, period, and wave profile analyses, were conducted using the “WaveAnalysis” Python script^6^. For medial sectional imaging, kymographs were generated and assessed using the same script, with modifications tailored to accommodate kymographs (these adaptations are now integrated into the script and detailed here: https://github.com/zacswider/waveAnalysis). In short, kymographs were vertically binned into lines approximately 4 microns wide, and the average pixel intensity across those 4 μ*m* was used to construct wave profiles. Subsequent analyses were the same as the previously reported procedures^6^. For wave profiles in Figure 1, the signal intensity was normalized to the maximum and minimum signal via

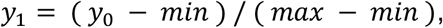

where y_0_ represents the raw signal, min is the minimum intensity, and max is the maximum intensity. The mean time-shifts illustrated in Figure S1E were produced utilizing a previously described MATLAB script^10^. Difference movies were utilized for all quantitative assessments. Figures were initially crafted in GraphPad Prism and subsequently refined and compiled in Adobe Illustrator. Statistical analyses were conducted using GraphPad Prism.

### Numeric simulation of ADSI and ADS models

Simulations of the ADSI model^4^ (Figures 2C,J) were performed on a square domain 200μ*m* × 200μ*m* with spatial resolution Δ*x* = 0.5 μ*m* and periodic boundary conditions for the earlier published set of parameters^4^ and *β* = 0.05. The simulations were initiated with random initial conditions spatially distributed in the vicinity of the uniform stationary state.

Simulations of the ADS model (Figures 2D,K) were performed on a square domain 100*μm* × 100*μm* with spatial resolution Δ*x* = 0.2 *μm* and periodic boundary conditions for the parameter values presented in Table S1 and *k*_2_ = 0.06 *M*^−2^*sec*^−1^. To induce wave dynamics, four spiral cores were embedded into the initial conditions.

Scatter plots (Figures 2H-K) were generated as follows. Fluorescent intensity signals of RhoA-GTP and total Rho (Figures 2H,I) were averaged within the boxes of 10x10 pixels. The resulting values were then normalized by the maximum and minimum values computed over all pixel boxes and all imaging frames of the respective imaging series as follows:

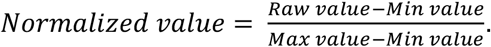

Simulation results (Figures 2J,K) for RhoA-GTP (RT) and total RhoA (TR, see Analysis of the ADS model below) were averaged inside the boxes of size 5x5 pixels and then normalized as above. Correlation coefficients (Figure 2H-K) were computed using the MathWorks MATLAB® function *corrcoef*.

### Analysis of the ADS model

We can quantitatively formulate a generic activator-depleted substrate model as a system of partial differential equations describing the spatio-temporal dynamics of the two membrane-associated forms of RhoA, GDP-bound inactive RD and GTP-bound active RT. Following earlier published work^9,43^ but assuming for simplicity that the cytoplasmic pool of inactive RhoA is so large that its concentration can be considered constant, we write:

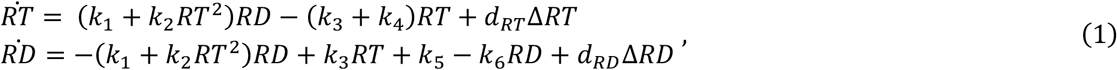

where the numbering of constants follows Figure S2A and their representative values are given in Table S1. In the following we omit the diffusion terms *d*_*RT*_ Δ*RT, d*_*RD*_ Δ*RD* and instead of *RD*, which cannot be assayed experimentally, we introduce the concentration of total RhoA GTPase *TR* = *RT* + *RD*, whose dynamics can be directly compared with that of IT-RhoA in experiments. This changes model (1) to the system of two ordinary differential equations:

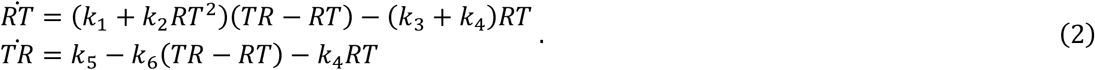

As a two-variable system of ordinary differential equations, model (2) can be fully characterized mathematically in terms of the analysis of its nullclines^19^. The nullclines of equations (2), obtained by solving 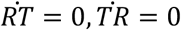, are then

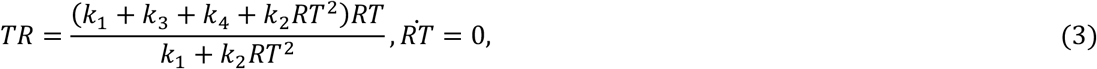

and

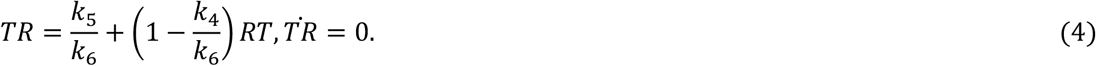

Existence of waves in the full spatially distributed model (1) requires that the nullclines (3,4) of the ordinary differential equations (2) satisfy the following conditions. First, nullcline (4) 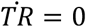, which is a straight line, must have a negative slope. Second, nonlinear nullcline (3) 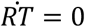 should have a descending segment with negative slope. An example of a phase portrait in which both conditions are satisfied is shown in Figure S2B. In this case, the point of intersection of the two nullclines corresponds to an unstable steady state of the type focus^19^ (empty circle, Figure S2B), which is surrounded by a stable limit cycle (green trajectory, Figure S2B).

These conditions impose strong restrictions on the model parameters. Indeed, for the nullcline 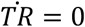 to have a negative slope, the rate of RT membrane detachment, *k*_4_, must be larger than that of the inactive RD, *k*_6_, (in practice, however, the requirement is even stronger, *k*_4_ ≫ *k*_6_). If the reaction of simultaneous inactivation and removal of the GTPase into the cytoplasm does not exist (*k*_4_ = 0 and the red arrow is removed from the diagrams in Figures 2B and S2A), nullcline (4), 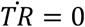, has a parameter-independent positive slope 1. In this case, under varying model parameters, the nullclines can intersect in one or three points. The model, therefore, could be either mono- or bistable, but cannot exhibit oscillations.

An important biological conclusion that immediately follows from the requirement

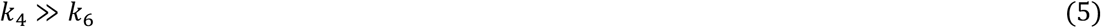

is that a generic activator-depleted substrate model (1) can explain cortical waves of GTPase activity only if the membrane residence time of active form of the GTPase is much shorter than that of its inactive form.

On the other hand, nullcline (3), 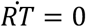, is a nonlinear function that has a sigmoidal shape with a descending segment only when *k*_1_ < (*k*_3_ + *k*_4_)⁄8, which can be readily seen from the analysis of the shape of nullcline (3). This strict inequality can be approximated as

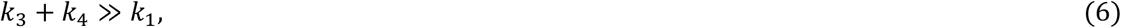

which could be biologically interpreted as a requirement that the rate of non-autocatalytic activation of a GTPase, *k*_1_, should be small in comparison with the total rate of GTPase inactivation, *k*_3_ + *k*_4_. In comparison with (5), this requirement is, however, much less restrictive.

Another important biological conclusion arises from the analysis of the direction of phase flow determined by the equations (2). This analysis shows (green arrows, Figure S2B) that the limit cycle, if it exists in the model at the given parameter values, is always traversed clockwise. In other words, this means that, contrary to the ADSI model, in the framework of the ADS model, the maximum of the total concentration of a GTPase always precedes that of the GTPase activity. Importantly, this conclusion is specified by the topological layout of the reaction network, it does not depend on the values of model parameters and is, thus, universal for the ADS model as defined by the network diagram in Figures 2B and S2A.

**Table S1.** The ADS model parameters.

|  |  |
| --- | --- |
| $k_1$ | $0.001 \text{ s}^{-1}$ |
| $k_2$ | $(0.05, 0.45) \text{ M}^{-2}\text{s}^{-1}$ |
| $k_3$ | $0.01 \text{ s}^{-1}$ |
| $k_4$ | $0.133333 \text{ s}^{-1}$ |
| $k_5$ | $0.0666667 \text{ M}^*\text{s}^{-1}$ |
| $k_6$ | $0.00444444 \text{ s}^{-1}$ |
| $d_{RT}$ | $0.05 \text{ }\mu\text{m}^2/\text{s}$ |
| $d_{RD}$ | $0.1 \text{ }\mu\text{m}^2/\text{s}$ |

