## Supplementary figures and images for "Testing models of cell cortex wave generation by Rho GTPases"

### Figure S1

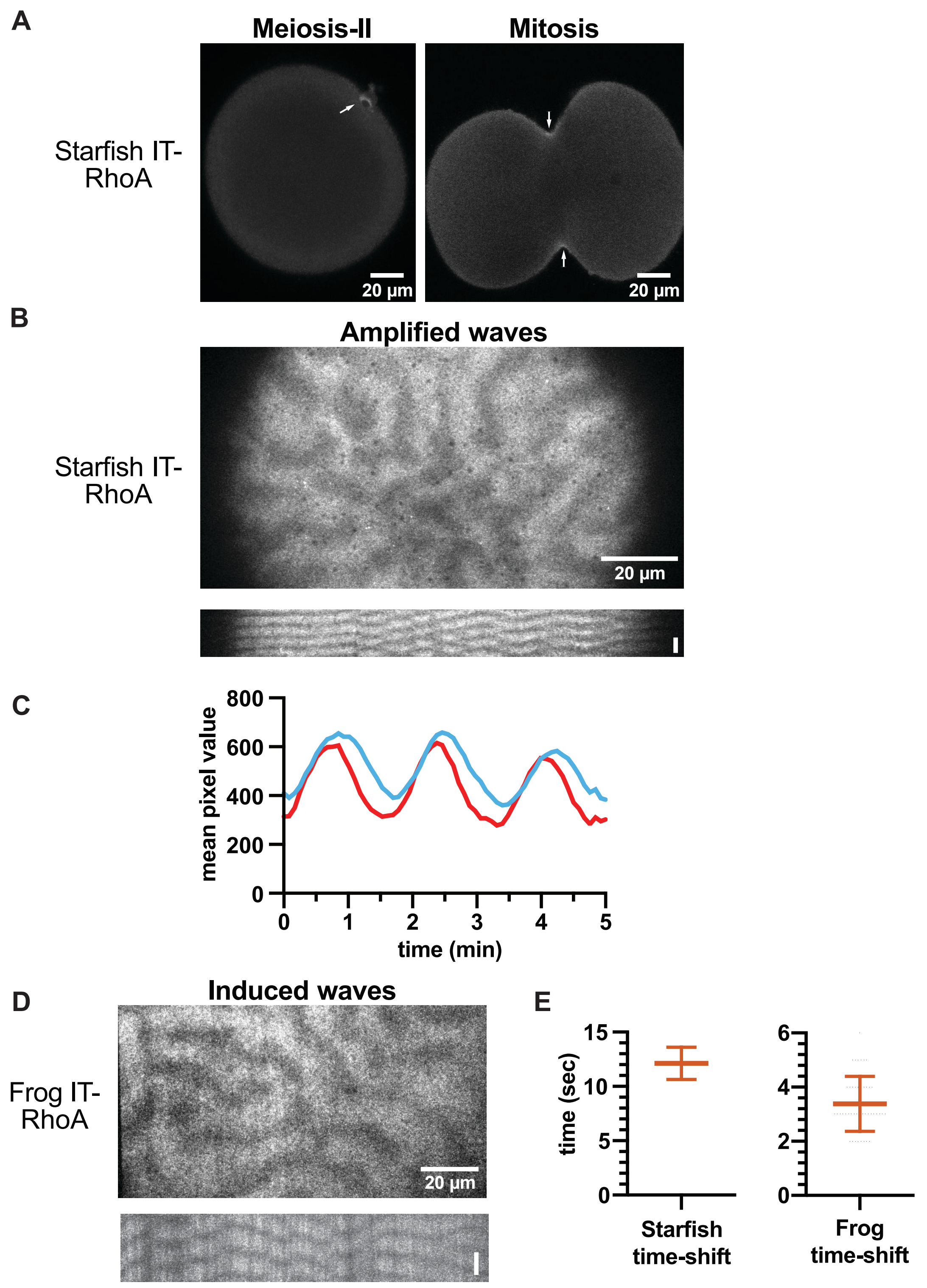

### Figure S2

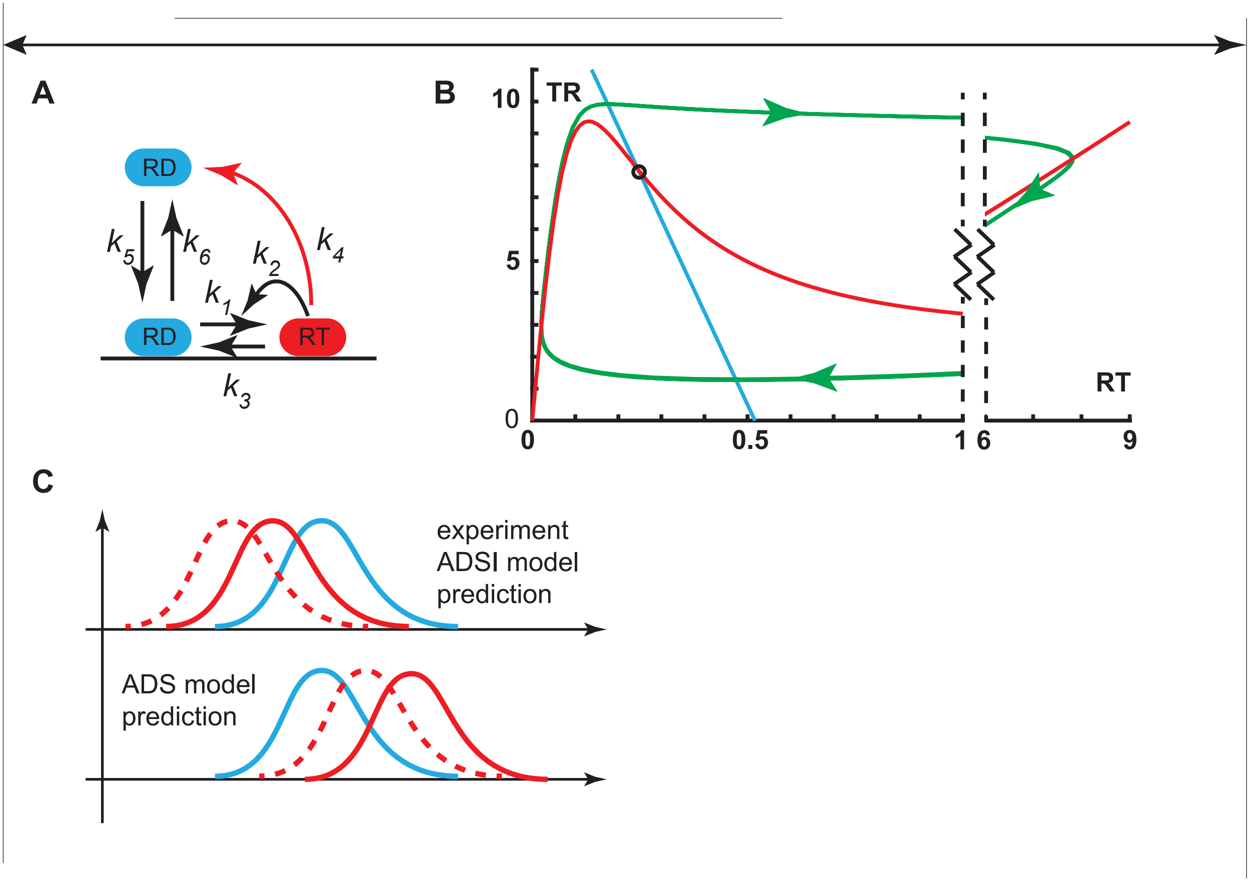
